# Mechanistic modeling of the SARS-CoV-2 disease map

**DOI:** 10.1101/2020.04.12.025577

**Authors:** Kinza Rian, Marina Esteban-Medina, Marta R. Hidalgo, Cankut Çubuk, Matias M. Falco, Carlos Loucera, Devrim Gunyel, Marek Ostaszewski, María Peña-Chilet, Joaquín Dopazo

## Abstract

Here we present a web interface that implements a comprehensive mechanistic model of the SARS-CoV-2 disease map in which the detailed activity of the human signaling circuits related to the viral infection and the different antiviral responses, including immune and inflammatory activities, can be inferred from gene expression experiments. Moreover, given to the mechanistic properties of the model, the effect of potential interventions, such as knock-downs, over-expression or drug effects (currently the system models the effect of more than 8000 DrugBank drugs) can be studied in specific conditions. By providing a holistic, systems biology approach to the understanding of the complexities of the viral infection process, this tool will become an important asset in the search for efficient antiviral treatments.

The tool is freely available at: http://hipathia.babelomics.org/covid19/

## Introduction

The recent pandemic of COVID-19 (Coronavirus Disease-2019), an emerging respiratory disease caused by the SARS-CoV-2 virus, which spread more efficiently than previous highly pathogenic coronaviruses SARS-CoV and MERS-CoV, has led to a tremendous toll of affected cases and over 56,000 fatalities in more than 180 countries since its first outbreak in late 2019 ^1^. Precisely due to the rapid transmission of this novel pathogen, no antiviral drugs or vaccines are available for SARS-CoV-2.

Understanding the molecular mechanisms that mediate SARS-CoV-2 infection is key for the rapid development of efficient preventive or therapeutic interventions against the COVID-19. A comprehensive description of such molecular mechanisms is represented in the corresponding disease map, that is, the fragment of the whole network of known human protein interactions that are relevant for the disease ^2^. The recent availability of a detailed catalog of viral-human protein interactions^3^ has facilitated the construction of a first version of the map of human molecular pathways involved in the viral infection. To do so, we firstly expanded the SARS-CoV-2 virus-human interactome from existing KEGG pathways maps to obtain a subset of signaling circuits (a circuit is the sub-pathway that defines the chain of signal transduction that connects a receptor protein to an effector protein) that represent a comprehensive model of the direct and indirect interactions of the virus with the cell. In particular, we selected those signaling circuits with at least one UniProt function that fit in one of these virus-related categories: 1) Host-virus interaction, 2) inflammatory response, 3) immune activity, 4) antiviral defense, 5) endocytosis. Our model is a part of a detailed repository of SARS-CoV-2 mechanisms, the COVID-19 Disease Map, constructed by an international community, and is available at: http://doi.org/10.17881/covid19-disease-map. Disease maps are repositories of knowledge of disease-relevant mechanisms that provide qualitative guidance for the interpretation of experimental findings. Actually, disease maps are the supporting foundation of different tools able to model the information contained in them in order to provide a detailed quantitative explanation for experimental results. In particular, mechanistic models of disease maps are becoming increasingly relevant for genomic data interpretation because they provide a natural link between omics data measurements and cell behavior and outcome, which ultimately accounts for the phenotype of the infection. The knowledge of these links allows a better understanding of the molecular mechanisms of the viral infection and the responses to drugs. Basically, mechanistic models map experimental values in the context of the disease map information, which is used to point out the relevant experimental, thus revealing aspects of the molecular mechanisms behind the experiment. It is important to note that this assessment is made from a systems biology perspective, in the holistic context of the disease map, and considers the functional interactions among the gene products as described in the map. Typically, these experimental values are gene expression transcriptomic data, although other data such as proteomic, phosphoproteomic, genomic, or even methylomics, can also be used.

However, beyond its usefulness for the functional interpretation of experimental results, the most interesting property of mechanistic models is that they can be used to predict the effects of interventions (inhibitions or over-activation, alone or in combinations) over proteins of the map in a given condition studied ^4^. Therefore, this opens the possibility of using these models for exploring new therapeutic options as well ^5^.

## Results

Here, we present the first implementation of a mechanistic model of the SARS-CoV-2 infection in a user-friendly interface. The model used here implements the *HiPathia* ^6^ algorithm, which has demonstrated to outperform other competing algorithms in a recent benchmarking ^7^. The mechanistic model implemented in *HiPathia* has been successfully used to understand the disease mechanisms behind different cancers ^6^ and was able to predict cancer vulnerabilities with a high precision ^8^. The model has been implemented in a user-friendly web application that inputs normalized gene expression values (or similar proteomics or phosphoproteomic values) and can be found at http://hipathia.babelomics.org/covid19/. As an example, we carried out some analyses that involve a case-control differential signaling analysis using a gene expression experiment with lung cell lines infected with SARS-CoV-2 (GEO id: GSE147507). The infected cells showed a differential activation pattern in circuits related to virus entrance to cell, activation of immune, inflammatory and other virus-triggered responses (see Figure 1A and Table 1 for a detailed list of differentially activated signaling circuits and Table 2 for detail on the differentially activated cell functionalities). Interestingly, several of the deregulated pathways include *TNF*, a target gene of chloroquine, one of the drugs with promising results against COVID-19 ^9^. Moreover, NF-kB signaling pathway has been highlighted in several studies as one of the main pathways responsible for COVID-19 progression ^10^ (Figure 1 B). Figure 1C depicts the heathmap of signaling activity profiles that discriminate the two classes of samples (cases and controls) compared. A very interesting option of the implementation of the model is the *Perturbation effect*. It allows estimating the effect of interventions (inhibitions or overexpression) across the signaling circuits of the model in a given condition. Moreover, the effect of more than 8000 targeted drugs from DrugBank can be predicted by selecting them, individually or in combinations. Figure 1D shows a simulated knockdown in protein *NFKB1A* and the cell functionality affected: *host-virus interaction*. Other options in the application allow building predictors or estimating the relevance of human mutations in the resulting infection phenotype.

**Figure 1.**
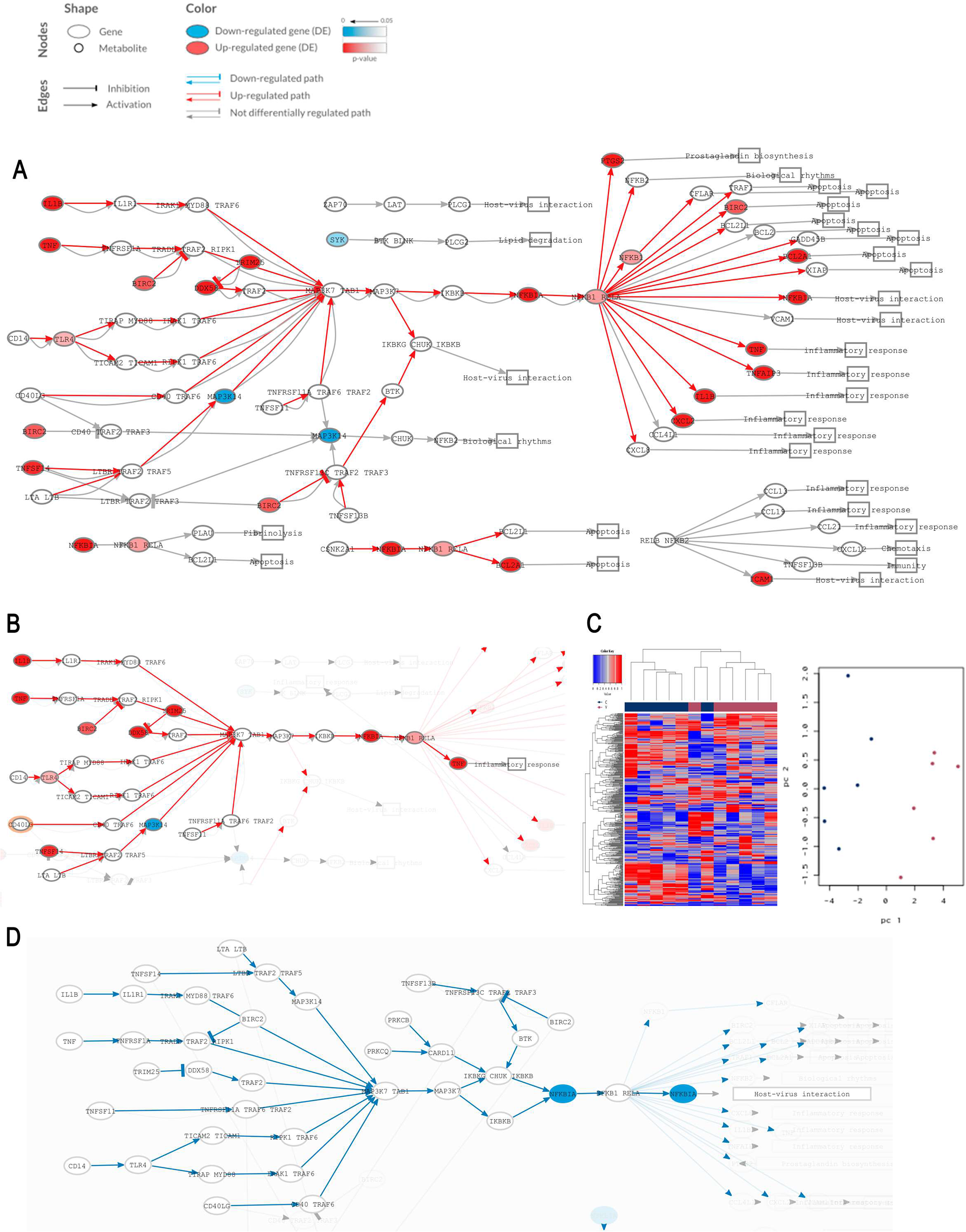
A) Activation pattern of NF-KB pathway in lung cell lines infected with SARS-CoV-2. B) Detail of NF-KB pathway’s circuit with TNF as effector protein. C) Heatmap representing activation values of all the circuits in COVID-19 disease map (left) and a representation of a Principal Component Analysis based on signaling profiles of the samples that clearly segregates the two conditions studied: controls are represented in dark blue and cases in purple (right). D) An example of the effect that in silico knockdown of gene NFKB1A (in blue) will have in the activation pattern of Toll-like signaling pathway in healthy lung tissue, using the Perturbation effect option of the software.

**Table 1:** Circuits from CoV-Hipathia differentially activated in lung cell lines infected with SARS-CoV-2.

| KEGG pathway: effector gene/s | UP/<br>DOWN | statistic | p-value<sup>1</sup> | FDR<sup>2</sup> | FC<sup>3</sup> | log FC |
| --- | --- | --- | --- | --- | --- | --- |
| MAPK signaling pathway: NLK | UP | 2.882 | 2.16E-03 | 0.023 | 1.098 | 0.135 |
| MAPK signaling pathway: STK3 | UP | 2.882 | 2.16E-03 | 0.023 | 1.209 | 0.274 |
| MAPK signaling pathway: ECSIT, TRAF6 | UP | 2.882 | 2.16E-03 | 0.023 | 1.182 | 0.241 |
| Ras signaling pathway: REL | UP | 2.882 | 2.16E-03 | 0.023 | 1.074 | 0.103 |
| Ras signaling pathway: MLLT4 | UP | 2.882 | 2.16E-03 | 0.023 | 1.049 | 0.069 |
| NF-kappa B signaling pathway: CFLAR | UP | 2.882 | 2.16E-03 | 0.023 | 1.212 | 0.278 |
| NF-kappa B signaling pathway: BIRC2 | UP | 2.882 | 2.16E-03 | 0.023 | 1.261 | 0.335 |
| NF-kappa B signaling pathway: XIAP | UP | 2.882 | 2.16E-03 | 0.023 | 1.176 | 0.234 |
| NF-kappa B signaling pathway: BCL2L1 | UP | 2.882 | 2.16E-03 | 0.023 | 1.193 | 0.254 |
| NF-kappa B signaling pathway: GADD45B | UP | 2.882 | 2.16E-03 | 0.023 | 1.179 | 0.238 |
| NF-kappa B signaling pathway: BCL2A1 | UP | 2.882 | 2.16E-03 | 0.023 | 1.570 | 0.651 |
| NF-kappa B signaling pathway: NFKB2 | UP | 2.882 | 2.16E-03 | 0.023 | 1.225 | 0.293 |
| NF-kappa B signaling pathway: CXCL8 | UP | 2.882 | 2.16E-03 | 0.023 | 1.185 | 0.245 |
| NF-kappa B signaling pathway: IL1B | UP | 2.882 | 2.16E-03 | 0.023 | 1.341 | 0.424 |
| NF-kappa B signaling pathway: TNFAIP3 | UP | 2.882 | 2.16E-03 | 0.023 | 1.311 | 0.390 |
| NF-kappa B signaling pathway: NFKBIA | UP | 2.882 | 2.16E-03 | 0.023 | 1.254 | 0.327 |
| NF-kappa B signaling pathway: PTGS2 | UP | 2.882 | 2.16E-03 | 0.023 | 1.267 | 0.341 |
| NF-kappa B signaling pathway: CXCL2 | UP | 2.882 | 2.16E-03 | 0.023 | 1.335 | 0.417 |
| NF-kappa B signaling pathway: IKBKG, CHUK, IKBKB | UP | 2.882 | 2.16E-03 | 0.023 | 1.110 | 0.151 |
| NF-kappa B signaling pathway: BCL2L1 | UP | 2.882 | 2.16E-03 | 0.023 | 1.098 | 0.135 |
| NF-kappa B signaling pathway: BCL2A1 | UP | 2.882 | 2.16E-03 | 0.023 | 1.448 | 0.534 |
| HIF-1 signaling pathway: TIMP1 | UP | 2.882 | 2.16E-03 | 0.023 | 1.051 | 0.072 |
| HIF-1 signaling pathway: TIMP1 | UP | 2.882 | 2.16E-03 | 0.023 | 1.051 | 0.072 |
| HIF-1 signaling pathway: EDN1 | UP | 2.882 | 2.16E-03 | 0.023 | 1.121 | 0.164 |
| HIF-1 signaling pathway: NOS2 | UP | 2.882 | 2.16E-03 | 0.023 | 1.585 | 0.664 |
| HIF-1 signaling pathway: PDK1 | UP | 2.882 | 2.16E-03 | 0.023 | 1.037 | 0.052 |
| HIF-1 signaling pathway: PGK1 | UP | 2.882 | 2.16E-03 | 0.023 | 1.048 | 0.068 |
| HIF-1 signaling pathway: LDHA | UP | 2.882 | 2.16E-03 | 0.023 | 1.047 | 0.066 |
| mTOR signaling pathway: VEGFA | UP | 2.882 | 2.16E-03 | 0.023 | 1.059 | 0.083 |
| mTOR signaling pathway: TSC1 | DOWN | -2.882 | 2.16E-03 | 0.023 | 0.946 | -0.080 |
| PI3K-Akt signaling pathway: BCL2L11 | DOWN | -2.882 | 2.16E-03 | 0.023 | 0.918 | -0.123 |
| Apoptosis: FADD, TRADD | UP | 2.882 | 2.16E-03 | 0.023 | 1.118 | 0.161 |
| Apoptosis: IRAK3, MYD88 | UP | 2.882 | 2.16E-03 | 0.023 | 1.167 | 0.223 |
| Toll-like receptor signaling pathway: CXCL10 | UP | 2.882 | 2.16E-03 | 0.023 | 1.518 | 0.602 |
| Toll-like receptor signaling pathway: IFNA1 | UP | 2.882 | 2.16E-03 | 0.023 | 1.486 | 0.572 |
| Toll-like receptor signaling pathway: IL1B | UP | 2.882 | 2.16E-03 | 0.023 | 1.172 | 0.229 |
| Toll-like receptor signaling pathway: IL6 | UP | 2.882 | 2.16E-03 | 0.023 | 1.526 | 0.610 |
| RIG-I-like receptor signaling pathway: MAPK14 | UP | 2.882 | 2.16E-03 | 0.023 | 1.214 | 0.280 |
| RIG-I-like receptor signaling pathway: MAVS, TMEM173 | UP | 2.882 | 2.16E-03 | 0.023 | 1.204 | 0.268 |
| RIG-I-like receptor signaling pathway: IRF7 | UP | 2.882 | 2.16E-03 | 0.023 | 1.540 | 0.623 |
| Ras signaling pathway: MAPK8 | UP | 2.722 | 4.33E-03 | 0.035 | 1.043 | 0.061 |
| NF-kappa B signaling pathway: TRAF1 | UP | 2.722 | 4.33E-03 | 0.035 | 1.193 | 0.254 |
| NF-kappa B signaling pathway: TNF | UP | 2.722 | 4.33E-03 | 0.035 | 1.639 | 0.713 |
| HIF-1 signaling pathway: TFRC | UP | 2.722 | 4.33E-03 | 0.035 | 1.049 | 0.068 |
| Sphingolipid signaling pathway: SMPD2 | UP | 2.722 | 4.33E-03 | 0.035 | 1.388 | 0.473 |
| Apoptosis: FADD, TRADD | UP | 2.722 | 4.33E-03 | 0.035 | 1.358 | 0.442 |
| Apoptosis: TRAF2, RIPK1, TRADD | UP | 2.722 | 4.33E-03 | 0.035 | 1.358 | 0.442 |
| Toll-like receptor signaling pathway: TNF | UP | 2.722 | 4.33E-03 | 0.035 | 1.434 | 0.520 |
| TNF signaling pathway: CASP7 | UP | 2.722 | 4.33E-03 | 0.035 | 1.388 | 0.473 |
| TNF signaling pathway: CASP3 | UP | 2.722 | 4.33E-03 | 0.035 | 1.371 | 0.455 |
| TNF signaling pathway: BAG4 | UP | 2.722 | 4.33E-03 | 0.035 | 1.391 | 0.476 |

P-value is calculated using Wilcoxon test.
2 FDR refers to p-value adjusted for multiple comparisons using Benjamini and Hochberg method.
3 FC means Fold Change.

**Table 2:** Functions from CoV-Hipathia differentially activated in lung cell lines infected with SARS-CoV-2.

| UniProt function | UP/DOWN | statistic | p-value<sup>1</sup> | FDR<sup>2</sup> | FC<sup>3</sup> | logFC |
| --- | --- | --- | --- | --- | --- | --- |
| Innate immunity | UP | 2.882 | 2.16E-03 | 0.018 | 1.048 | 0.067 |
| Immunity | UP | 2.882 | 2.16E-03 | 0.018 | 1.019 | 0.028 |
| Inflammatory response | UP | 2.882 | 2.16E-03 | 0.018 | 1.028 | 0.041 |
| Antiviral defense | UP | 2.882 | 2.16E-03 | 0.018 | 1.192 | 0.253 |
| Pyrogen | UP | 2.882 | 2.16E-03 | 0.018 | 1.177 | 0.235 |
| Prostaglandin biosynthesis | UP | 2.882 | 2.16E-03 | 0.018 | 1.267 | 0.341 |
| Prostaglandin metabolism | UP | 2.882 | 2.16E-03 | 0.018 | 1.267 | 0.341 |
| Fatty acid biosynthesis | UP | 2.882 | 2.16E-03 | 0.018 | 1.266 | 0.341 |
| Lipid biosynthesis | UP | 2.882 | 2.16E-03 | 0.018 | 1.266 | 0.341 |
| Acute phase | UP | 2.882 | 2.16E-03 | 0.018 | 1.526 | 0.610 |
| Osteogenesis | UP | 2.882 | 2.16E-03 | 0.018 | 1.181 | 0.241 |
| Autophagy | DOWN | -2.722 | 4.33E-03 | 0.028 | 0.976 | -0.035 |
| Translation regulation | UP | 2.722 | 4.33E-03 | 0.028 | 1.015 | 0.022 |
| Sphingolipid metabolism | UP | 2.722 | 4.33E-03 | 0.028 | 1.388 | 0.473 |
| Necrosis | UP | 2.562 | 8.66E-03 | 0.046 | 1.009 | 0.013 |
| Fibrinolysis | UP | 2.562 | 8.66E-03 | 0.046 | 1.103 | 0.141 |
| Plasminogen activation | UP | 2.562 | 8.66E-03 | 0.046 | 1.072 | 0.101 |

P-value is calculated using Wilcoxon test.
2 FDR refers to p-value adjusted for multiple comparisons using Benjamini and Hochberg method.
3 FC means Fold Change.

Despite the limitations due to the few samples available, the results of the example clearly show the usefulness of this tool for modelling the repertoire of cell responses triggered by SARS-CoV-2, and the enormous potential that it has for future COVID-19 research and discovery of therapeutic interventions.

## Acknowledgements

This work is supported by grants SAF2017-88908-R from the Spanish Ministry of Economy and Competitiveness and “Plataforma de Recursos Biomoleculares y Bioinformáticos” PT17/0009/0006 from the ISCIII, both co-funded with European Regional Development Funds (ERDF) as well as H2020 Programme of the European Union grants Marie Curie Innovative Training Network “Machine Learning Frontiers in Precision Medicine” (MLFPM) (GA 813533) and “ELIXIR-EXCELERATE fast-track ELIXIR implementation and drive early user exploitation across the life sciences” (GA 676559).

